# Autophagy shapes the peptide repertoire of rheumatoid arthritis-associated HLA class II alleles

**DOI:** 10.64898/2026.03.24.713950

**Authors:** Natacha Madelon, Michael Stumpe, Julien Racle, Mégane Pluess, David Cune, Alessandra Notto, Sebastien Viatte, Essia Saiji, Nataliya Yeremenko, Jakob Nilsson, David Gfeller, Caroline Ospelt, Joern Dengjel, Monique Gannagé

**Affiliations:** Service of Immunology and Allergy, Lausanne University Hospital, University of Lausanne, 1011 Lausanne, Switzerland; Department of Biology, University of Fribourg, Chemin du Musée 10, 1700 Fribourg Switzerland; Department of Oncology, Ludwig Institute for Cancer Research, University of Lausanne, Lausanne, Switzerland; Agora Cancer Research Center, Lausanne, Switzerland; Swiss Cancer Center Leman (SCCL), Geneva, Switzerland; Swiss Institute of Bioinformatics (SIB), Lausanne, Switzerland; Lydia Becker Institute of Immunology and Inflammation, Faculty of Biology, Medicine and Health, The University of Manchester; NIHR Manchester Musculoskeletal Biomedical Research Centre, Central Manchester NHS Foundation Trust, Manchester Academic Health Science Centre, Manchester, United Kingdom; Department of Pathology and Immunology, Faculty of Medicine, University of Geneva; Department of Clinical Immunology and Rheumatology and Department of Experimental Immunology, Academic Medical Center/University of Amsterdam, Amsterdam, the Netherlands; Department of Immunology, University Hospital of Zurich, Zurich, Switzerland; Department of Rheumatology, Center of Experimental Rheumatology, University Hospital of Zurich, University of Zurich, Zurich, Switzerland

**Author notes:** Correspondence to: Monique Gannagé. equal contribution (Natacha Madelon, Michael Stumpe).

## Abstract

Alternative pathways of antigen presentation are crucial in different immunological contexts such as autoimmunity and anti-microbial defense. Among these pathways, autophagy has a central role in delivering cytosolic substrates to the MHC class II compartment. However, its contribution to endogenous MHC class II presentation was only demonstrated for a few antigens. Here we focused our study on the contribution of autophagy to the peptidome of one major allele of the HLA-DR shared epitope, HLA DRB1*04:01 conferring the greatest risk factor for the development of rheumatoid arthritis (RA). We provide an extensive qualitative and quantitative mass spectrometry analysis of the autophagy related MHC class II peptide repertoire of the human DRB1*04:01 allele. A fraction of peptides representing 30% of the repertoire differ profoundly between autophagy sufficient and deficient cells. Our analysis demonstrates that autophagy contributes to MHC class II presentation of peptides from seven described RA autoantigens, the majority of them being related to the ER folding and stress response (calreticulin, calnexin, the 78 kDa glucose-regulated protein (GRP78)-also known as binding immunoglobulin protein (BiP) and several protein from the heat-shock-protein 70 family). Our results correlate with an increased activation of autophagy, *in situ,* in synovial biopsies and synovial fibroblast (SF) of RA patients. We could further show that SF upregulate MHC class II and present peptides from autophagy related auto-antigens to CD4 T cells from RA patients. Our finding identifies autophagy as a potential process that could contribute to the break of peripheral tolerance during RA.

## Introduction

Endogenous alternative pathways of MHC class II presentation play an important role in shaping the CD4 T cell response, in different contexts such as central and peripheral tolerance, or in the adaptive immune response to tumors or pathogens. These pathways have regained interest not only in non-professional antigen presenting cells (APCs) that lack phagocytic capacity, but also in professional APCs.

In professional APCs, the contribution of endogenous pathways to MHC class II presentation has been overlooked although there is some evidence that they could contribute to the MHC class II peptidome and therefore impact the priming phase of CD4 T cell responses. Indeed, in dendritic cells (DCs) an important proportion of naturally presented peptides (from 20 to 50%) are derived from cytosolic and nuclear proteins ^1–3^. Since DCs contribute to both central and peripheral tolerance mechanisms, understanding the pathways implicated in the generation of their steady state ligandome is of potential interest.

In non-professional APCs, which can upregulate MHC class II expression upon inflammation, sampling the intracellular milieu in the context of infection ^4^, autoimmunity, antitumoral responses ^5^ or alloreactivity ^6^ can impact the effector phase of the CD4 T cell response. In this respect different non-hematopoietic cells such as endothelial cells, epithelial cells, tumor cells, or fibroblasts have been described to upregulate MHC Class II molecules and contribute to the adaptive immune response in several pathological settings. In the tumor microenvironment in particular, numerous studies have shown that MHC Class II expression by cancer associated fibroblasts, by lymph node stromal cells or by the tumor cells directly contribute to the antitumoral response in different settings ^7^.

In the context of autoimmunity, fewer studies have demonstrated the role of MHC Class II expression in non-professional APCs, such as epithelial and endothelial cells. Indeed, while MHC Class II expression by thymic epithelial cells is well established and demonstrated to be essential for central tolerance and CD4 T cell selection, the contribution of MHC class II expression in non-professional APCs to peripheral tolerance mechanisms is less studied and still controversial. Among these cells, MHC class II expression in lymph node stromal cells ^8^, was shown to promote peripheral tolerance and support Tregs expansion and function in age related autoimmune disorders. More recently several studies have focused on the role of MHC Class II expression in intestinal epithelial cells ^6,9^.

Since endogenous antigen presentation is the prevalent mechanism of antigen presentation in these non-canonical APCs, it is relevant to dissect in these cells the intracellular pathways responsible for MHC class II presentation. Autophagy is the major pathway responsible for delivering intracellular antigens to the MHC class II compartment. The contribution of autophagy in thymic epithelial cells to central tolerance mechanisms has been demonstrated in mouse models ^10,11^. However, while some indirect evidence for the implication of autophagy in autoimmune disorders exists, there is a lack of direct demonstration of the contribution of the pathway to MHC class II presentation of auto-antigens and its impact on peripheral tolerance mechanisms.

Here we aimed to analyze the autophagy-related MHC class II peptidome of the shared epitope that confers a high risk of developing rheumatoid arthritis. We show that autophagy impacts the MHC class II peptide repertoire of the DRB1*04:01 allele, both qualitatively and quantitatively. As a consequence, in the context of rheumatoid arthritis (RA), and DRB1*04:01 restriction, MHC class II presentation of peptides from seven autoantigens described in the context of RA is significantly reduced when autophagy is compromised. These data correlate with increased activation of autophagy in synovial biopsies and synovial fibroblasts of RA patients. By peptide profiling of DR4 synovial fibroblasts, which are key players in the pathogenesis of the disease, we demonstrate that some of these epitopes are presented by synovial fibroblasts, which are unconventional APCs. Interestingly, upregulation of autophagy in synovial fibroblasts, significantly enhances activation of CD4 T cells from RA patients. Our study highlights a contribution of autophagy to MHC class II presentation of RA-related autoantigens and its potential role in the pathogenesis of this autoimmune disease.

## Results

### Autophagy contributes to the processing of DR4 restricted autoantigens

We focused our analysis on a DRB1*04:01 mono-allelic restricted repertoire. Indeed, mass spectrometry analysis of a mono-allelic repertoire is more informative for the purpose of our study, by circumventing bias due to differences in peptide-binding rules and by providing a more scalable experimental set up. DRB1*04:01 is associated with rheumatoid arthritis (RA) as being an allele from the shared epitope. Thus, this analysis could highlight if HLA presentation of epitopes from RA-related autoantigens could be autophagy dependent. We used a 293T cell line engineered to constitutively express DRB1*04:01, and deleted Atg12, an autophagy essential gene, by CRISPR-Cas9 technology **(Fig. S1A).** We verified that the level of HLA-DR4 was not affected by autophagy deletion **(Fig. S1B).** We then analyzed the peptides eluates of MHC class II molecules by mass spectrometry (MS). For all experiments, both eluates from wild-type and autophagy-deficient cells were paired and analyzed at the same time **(Fig. 1A).** To determine peptides that were presented by HLA-DRB1*04:01 in the MHC-II peptidomics data, we used MixMHC2pred-2.0 ^12^. Peptides were retained as HLA-DRB1*04:01 ligands when their % Rank returned by MixMHC2pred was below 20, corresponding to a threshold of predicted weak binding. Peptides of length shorter than 12 amino acids or longer than 21 amino acids were not considered as these are not predicted by MixMHC2pred and are mostly contaminants. To compare the expression of peptides between the two genotypes, peptides from the same source proteins were compared across five experiments, after normalizing the intensity of each peptide per run. We found that the length and the motifs of peptides eluates were not different between both genotypes, as the classical main DRB1*04:01 motifs was predominant in the two cell lines (as analysed by MoDec^13^) **(Fig. S1C).** However, qualitative and quantitative changes were observed as we found that an average of 24% of peptides (range16-29%, n=5) from the DRB1*04:01 restricted repertoire were significantly enriched in Atg12^+/+^ cells compared to the Atg12^−/-^ cells, **(Fig. 1B; and Supplemental Table 1 and Table 2).** In parallel but to the same extent a proportion of peptides, with an average of 31% (range 29-33%, n=5) from Atg12^−/-^ cells were also enriched in the absence of autophagy **(Fig. S1D),** arguing for the fact that alternative MHC Class II endogenous pathways, such as chaperon-mediated autophagy could be activated in the absence of autophagy. We then analyzed the cellular localization of source proteins for peptides enriched in autophagy sufficient cells. Gene ontology (GO) analysis revealed that most enriched peptides in autophagy sufficient cells originated from proteins of the intracellular compartment, with significant enrichment of pathways related to protein refolding, antigen processing, protein folding in endoplasmic reticulum and the unfolded protein response (**Fig. 1C)**. This was reflected by enriched MHC-II presentation of several peptides from heat shock proteins and chaperones in autophagy-competent cells **(Fig. 1D).** These proteins (n=17) were mainly from the cytoplasm, its organelles, and their membrane-associated proteins. Interestingly, half of these proteins were known autoantigens **(Fig. 1E)** linked to various autoimmune conditions. In particular, in the context of RA, six of these auto-antigens were reported to elicit either a T cell response or/and more often an autoantibody response: Calreticulin (CALR), Calnexin (CANX), GRP78 (78 KDa glucose-regulated protein) also known as BIP/HSPA5, several Heat shock 70KDa proteins, Glyceraldehyde-3-dehydrogenase (GAPDH), and the Fatty acid synthase (FASN). In contrast, we could not find any known autoantigens among proteins enriched in autophagy-deficient cells. However, and interestingly, the most enriched peptides, in autophagy-deficient cells were derived from MHC class I molecules **(Fig. S1E and F)**, in accordance with data from our group and from others^14^, describing autophagy as responsible for MHC class 1 recycling, degradation and, ER quality control ^15^.

**Figure 1.**
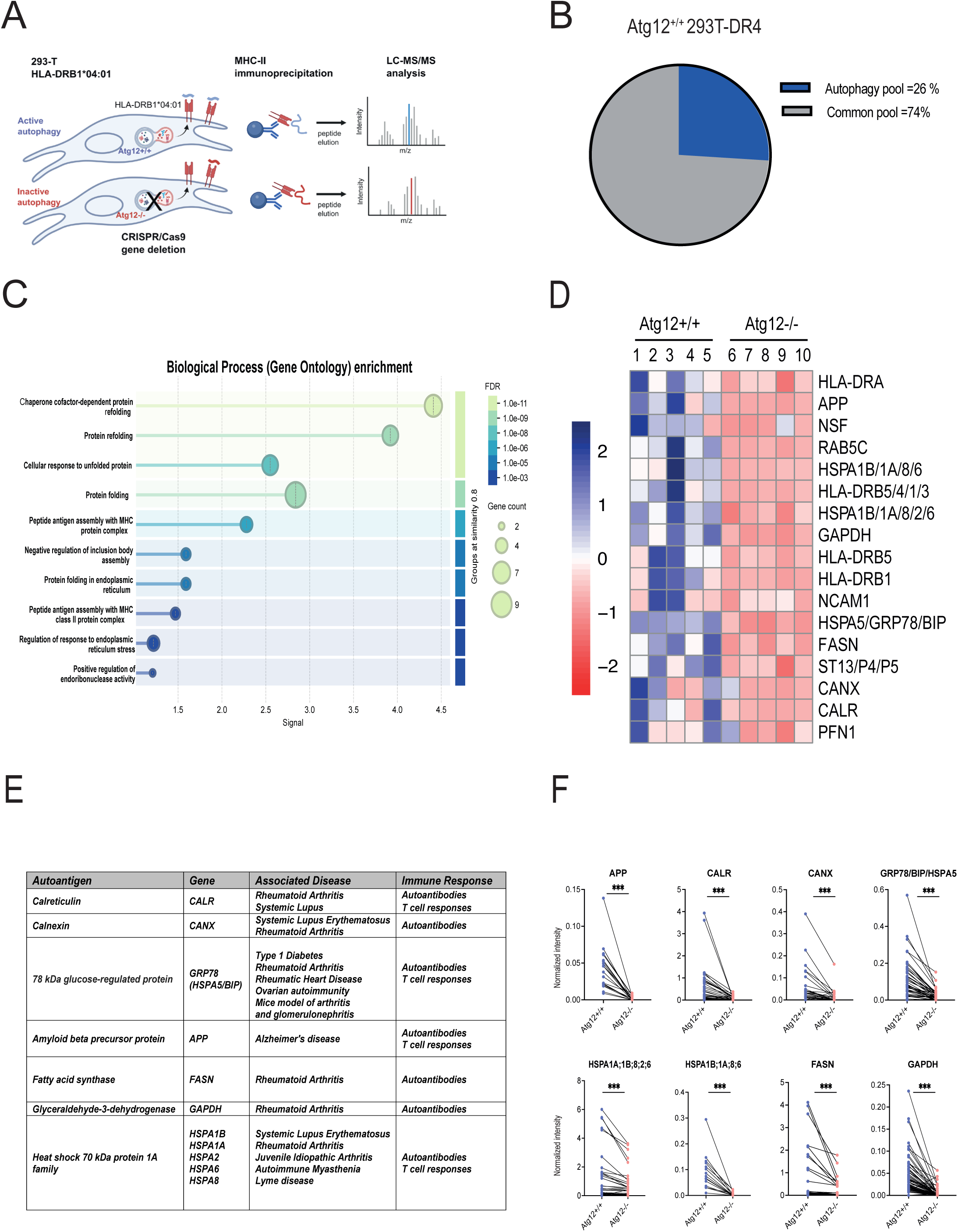
Autophagy contributes to the processing of DR4 restricted autoantigens. **(A)** Figure showing the experimental design: MHC Class II molecules were immunoprecipitated from HLA-DRB1*04:01 cells that are either autophagy sufficient (Atg12+/+) or deficient (Atg12-/-), and analyzed simultaneously by mass spectrometry. **(B)** Pie chart showing the number and relative abundance (%) of peptides that are enriched (dark blue) in the MHC class II peptidome of autophagy sufficient (Atg12^+/+^) 293T-DR4 cells as compared to (Atg12^−/-^) paired sample. The gray repertoire represents the pool of peptides that are stable between (Atg12^−/-^) and (Atg12^+/+^) 293T-DR4 cells. One representative paired experiment is shown out of 5. **(C)** GO Biological Process enrichment analyzis is shown (STRING software) on the indicated gene set. Significantly enriched GO terms are clustered based on semantic similarity (similarity threshold = 0.8). Node size represents the number of genes associated with each GO term, and node color indicates the false discovery rate (FDR) of enrichment, as shown in the scale. Edges denote similarity relationships between GO terms. **(D)** Heat Map representing the relative intensity of all proteins that are significantly enriched in MHC Class II peptides eluates from Atg12^+/+^ vs Atg12^−/-^ -DRB1*04:01 cells. A z normalization of the MS intensity was done for every antigen. **(E)** Table showing autoantigens that are statistically enriched in the peptidome of autophagy sufficient DRB1*04:01 cells**. (F)** Graphs showing the normalized intensity (in % of the repertoire) of every peptide from the related enriched autoantigen.

The selective enrichment of several native autoantigens in autophagy-sufficient cells highlights autophagy as an important endogenous pathway for MHC Class II processing of autoantigen related epitopes, that originate from the intracellular compartment. To get a quantitative estimate of the occupancy of the repertoire for every given autoantigen, we summed the relative frequencies of their related peptides. Accordingly, and interestingly, by analyzing their overall relative abundance within the repertoire **(Fig. 1F),** we could estimate that autoantigens described to elicit both a T cell and a B cell response in RA patients (HSP70^16,17^, GRP78^18^ and CALR^19,20^ were statistically overrepresented in the autophagy related peptidome and constitute around 21% of the MHC class II repertoire of DR4 cells. Among these autoantigens, the most represented ones’ in the peptidome were peptides from HSP70 protein 1A family (9.5% of the repertoire), from Calreticulin (around 4.4% of the peptide repertoire) followed by GRP78 (1% of the repertoire). For the 3 remaining autoantigens (GAPDH, CANX, FASN) described to elicit only an autoantibody response in RA patients^21^, the fatty acid synthase (FASN) was the most represented one, with some epitope representing around 4% of the MHC class II peptide repertoire of DR4 cells, while the two other remaining autoantigens represented around 0.9%. Overall peptides from known RA autoantigens that are enriched in autophagy-sufficient cells represent around 21% (range between 13.7-26.1% in five independent experiments) of the peptide repertoire of the DR4 allele.

### Increased activation of autophagy in synovial fibroblasts of RA patients

In order to address our finding in the context of the physiopathology of the disease, we first analyzed the transcriptional expression of autophagy essential genes in 23 synovial biopsies of RA patients, in an active phase of their disease. We used as a control a cohort of 20 osteoarthritis (OA) patients, and 8 healthy donors (HD). By Q-PCR analysis, at least 3 mRNA transcripts of essential autophagy genes were significantly upregulated (Atg8/LC3, Atg5 and Atg12) in both RA and OA patients **(Fig. 2A).** Because mRNA transcripts upregulation does not necessarily corelate with an increase in autophagosomes formation and is not a unique hallmark of autophagy upregulation, we evaluated in parallel, by immunofluorescence analysis, the number of autophagosomes in 10 sections of synovial biopsies of both RA and OA patients. We found a significant increase in the number of autophagosomes in biopsies of RA patients as compared to OA patients **(Fig.2B),** that does not correlate with an increase in P62/SQTM1 staining **(Fig. S2A).** We therefore corelate the accumulation of autophagosomes in RA patients with an increase in autophagy flux. Because synovial fibroblasts (SF) are central to the pathogenesis of the disease, we analyzed the accumulation of autophagosomes in SFs. We found that the number of autophagosomes in SFs *in situ* (vimentin positive cells) was higher compared to other cell types **(Fig 2C)** an observation that was already reported by others ^22^.

**Figure 2.**
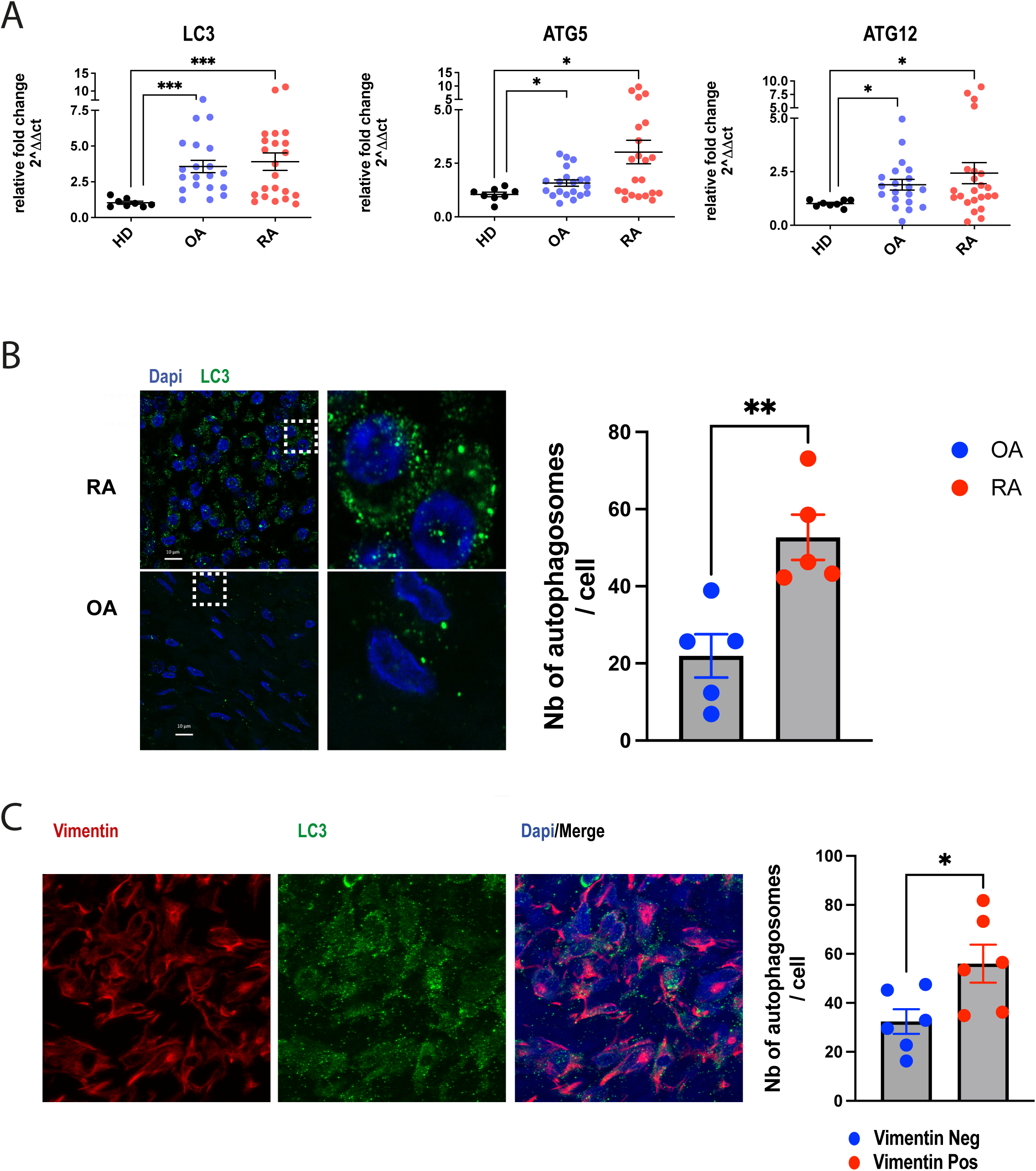
Autophagy is upregulated in the synovium of rheumatoid arthritis patients. **(A)** Quantitative PCR analysis of autophagy genes (LC3, Atg5 and Atg12) in mRNA extracts from synovial biopsies of rheumatoid arthritis patients (RA, n=23), osteoarthritis patients (OA, n=20) and healthy controls (HD, n=8). Graphs represent the relative fold change calculated as compared to the housekeeping gene (GAPDH). **(B)** Immunofluorescence analysis of autophagosomes accumulation in paraffin-embedded sections from synovial biopsies of 5 OA and 5 RA patients. Left: one representative staining of LC3 is shown (LC3 in green, DAPI in blue). Scale bar is in white. Right: Bar graph showing the mean number of autophagosomes/cell per patient. For each synovial biopsy, confocal analysis of Z stacks for at least 3 field were computed with a total of at least 150 cells per patient. Images were analyzed using Imaris software, autophagosomes were computed as LC3 positive vesicles from 200 to 500 nm. **(C)** Immunofluorescence analysis of autophagosomes accumulation in synovial fibroblasts from paraffin-embedded sections of synovial biopsies of RA patients. One representative section is shown. Autophagosomes: LC3+ vesicles in green, Synovial fibroblasts (SF): vimentin+ cells in red. 6 RA patients were analyzed, the mean number of autophagosomes per vimentin+ cell is 56 (SD=19) and the mean number of autophagosomes per vimentin^neg^ cell is 32 (SD=12)

### Synovial fibroblasts of RA patients upregulate autophagy and activate autologous CD4 T cells

We hypothesized that SFs by upregulating autophagy could serve as non-canonical antigen presenting cells under inflammatory conditions. Indeed, among their role in the pathogenesis of RA, SF have been described to interact with autoreactive CD4 T cells contributing to their activation^23^. Interestingly, in synovial biopsies of RA patients, co-staining analysis showed that SF upregulate MHC class II expression *in situ* **(Fig. 3A),** during RA, as reported by other studies ^24^. In parallel, *in vitro,* we could upregulate MHC class II expression in primary synovial fibroblasts after interferon-gamma (IFNg) treatment **(Fig. 3B).** Since SF are key cells in the pathogenesis of RA, we decided to evaluate the MHC class II peptidome of these cells, and the potential contribution of autophagy to their peptidome. We therefore analyzed the MHC class II peptide repertoire of 3 SF primary lines derived from 3 RA patients (SKH100, SKH124 and SKH108). Two of them being HLA-DRB1*0401 (SKH100, SKH108) and one serving as a negative control (SKH124). This evaluation was of particular interest, since the MHC class II peptidome of SF is not yet characterized. Interestingly, from the SF peptidome analysis **(Supplemental Table 3),** we identified DRB1*04:01 restricted peptides from 13 different RA related auto-antigens, and among them, we identified GAPDH, CALR, FASN, and Hsp70 proteins that we had characterized in the first section of this study to be autophagy dependent **(Fig. 3C-E).** These peptides were relatively abundant since their frequency in the MHC class II peptidome of SF was over 1%. Given the fact that autophagosomes accumulate in SF during the course of RA, autophagy could participate to the processing of autoantigens in SF, in particular to their role as non-canonical antigen presenting cells, and therefore activate pathogenic autoreactive CD4 T cells. In order to demonstrate that, we cocultured PBMCs from 4 RA patients with their matched autologous SF that were previously starved in order to upregulate autophagy, and cultured with IFNg in order to upregulate MHC class II **(Fig. S2B-D).** In the condition where autophagy was upregulated upon starvation in SF, autologous T cells were more activated and secreted more IFNg, arguing for a contribution of autophagy in autoantigen presentation in SF **(Fig. 3F).** Therefore, autophagy upregulation in RA SF is a potential mechanism implicated in the pathogenesis of the disease. Indeed, autophagy by contributing to MHC-II presentation of several RA autoantigens in synovial fibroblasts that are key players in the disease, could contribute to autoimmunity.

**Figure 3.**
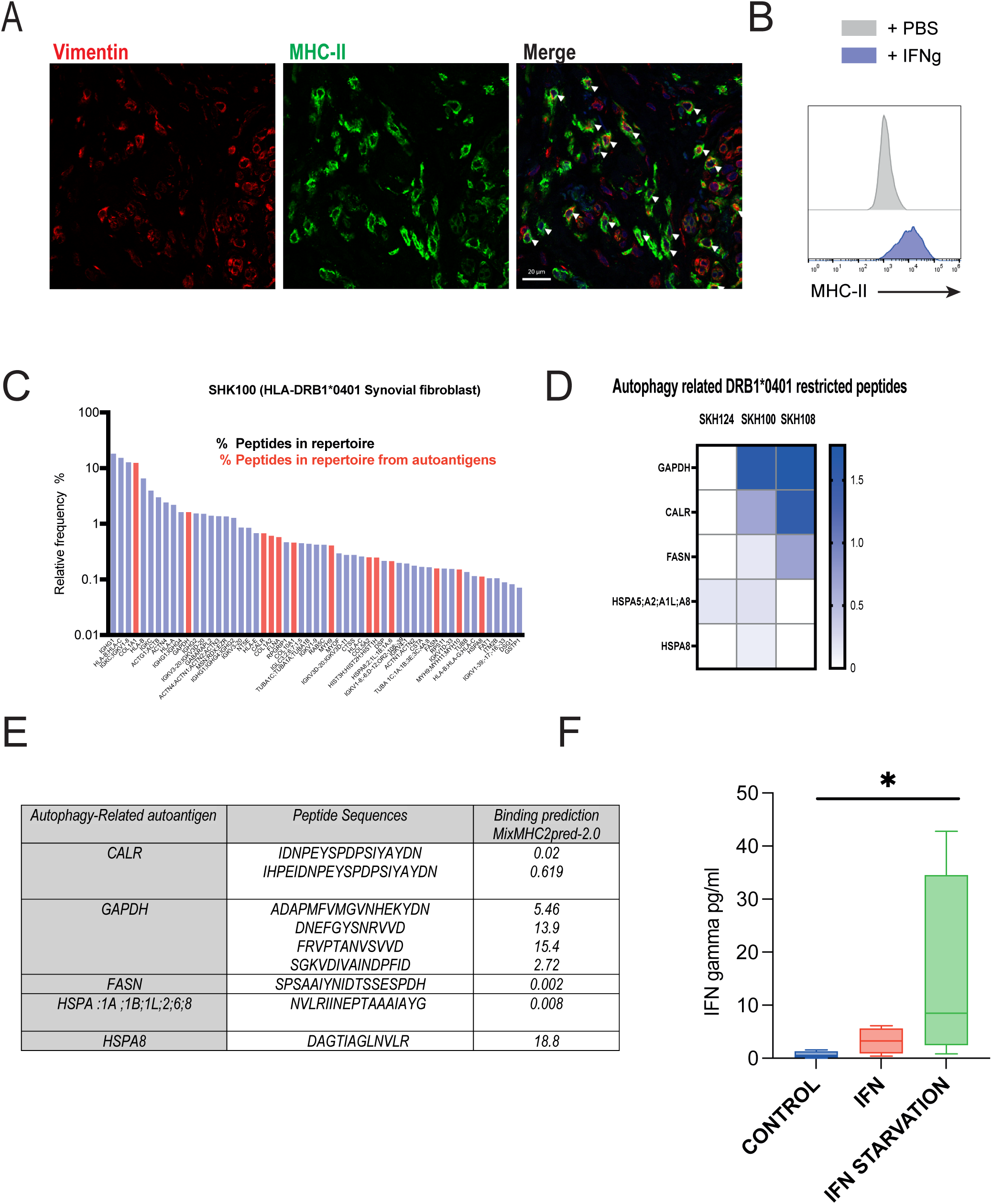
Synovial fibroblast of RA patients upregulate autophagy and activate autologous CD4 T cells. **(A)** Immunofluorescence analysis of MHC class II expression in synovial fibroblasts in paraffin-embedded section from synovial biopsies of RA patients. One representative staining is shown (vimentin in red, MHC class II in green, DAPI in blue). White arrows indicate MHC class II staining in vimentin positive cells. A total of 4 RA patients were analyzed, 25% of vimentin+ cells are MHC class II positive. **(B)** Histogram showing MHC Class II expression in synovial fibroblast after 48 hours of INF gamma treatment. **(C)** Graph showing the profile of Mass Spectrometry (MS) relative intensity of peptides eluates from MHC class II molecules of SHK100 SF primary line from an HLA-DRB1*04:01 RA patient. Peptides that are known autoantigens are depicted in red. The Y axis represents the normalized intensity and the X axis the related source proteins. MS intensities of every peptide were summed for every given protein and normalized to the sum of MS intensities of the run. **(D)** Heat map representing the relative frequency of autophagy related peptides in the MHC Class II peptidome from 3 SF primary lines (SKH124 a control non HLA-DR4 line and SKH108 and SKH100, two HLA-DR4 lines) **(E)** Table showing peptides sequences from autophagy related peptides that are present in the peptidome of synovial fibroblasts, and their binding prediction to HLA-DRB1*4:01 allele **(F)** Bar graphs represent IFN-gamma secretion assay from matched co-cultures of PBMCs from RA donors (n=3) with their autologous synovial fibroblast. Synovial fibroblasts are pre-cultured with IFNg for 48h, in normal complete medium or in starvation medium in order to upregulate autophagy.

## Discussion

The contribution of autophagy to autoimmune and inflammatory disorders have been suggested by different genome-wide association studies (GWAS) ^25^. In that context, the role of autophagy in Crohn disease was clearly established, with the identification of an ATG16L1 variant associated with the disease and responsible for a functional defect of the pathway ^26^ leading to a decrease in bacterial clearance and an increase in IL-1 beta production. During lupus, a SNP in the *ATG5* gene ^27^ seems also to be associated with disease. In RA, a SNP in the PRDM1-ATG5 region has been described to be associated with the disease ^28^. In parallel, different studies have demonstrated the upregulation of autophagy in both synovial fibroblasts and monocytes during the course of the disease. In particular, a role of the pathway in SF resistance to apoptotic cell death was demonstrated ^22^. Interestingly, TNF alpha -a key cytokine in the pathogenesis of RA-contributes to the upregulation of autophagy in these cells, and to an increase in their ER stress ^29^.

Importantly, autophagy seems to control citrullination that contributes to the break of tolerance, and the formation of anti-citrullinated protein antibodies (ACPA) during RA ^30^. Indeed, in both mouse and human models, activation of PAD enzymes requires autophagy, resulting in the citrullination of selected autoantigens such as filaggrin ^31^, alpha-enolase and vimentin ^32,33^. The precise mechanism linking the pathway to PAD activity is not yet characterized. While these studies have pointed towards a contribution of autophagy to the pathogenesis of autoimmune disorders, none has explored the role of the pathway in MHC-II presentation of native auto-antigens.

Our study of a monoallelic DRB1*04:01 repertoire from the shared epitope has shown that several described autoantigens are delivered to the MHC-II compartment by autophagosomes. Among them, some RA related autoantigens were enriched in autophagy sufficient cells and have been shown to elicit either a T cell response or an antibody response in RA patients. Among these antigens, HLA-DR restricted peptides from calreticulin, HSPA1A and HSPA5 (also known as BIP or GRP78) were identified in synovial tissue of RA patients ^17,34^. In parallel, proteomic analysis of synovial biopsies identified 11 proteins that strongly corelate with the inflammatory clinical score, among them HSPA1A family proteins, Calreticulin and BIP/GRP78 ^35^. Interestingly, HSPA1A family proteins, Calreticulin and BIP/GRP78 are proteins that are involved in the ER stress a pathway, that is activated during the inflammatory phase of RA. Moreover, calreticulin was shown to directly bind to GRP78/HSPA5. The calreticulin peptide (300-315), covering around 4% of the DR4 repertoire, that we have identified to be autophagy dependent, was already described to be one of the most abundant peptide of the DR4 peptidome ^36^. Interestingly calreticulin was demonstrated to be a selective substrate of autophagosomes ^37^, and a direct interactor of LC3 with an identified LIR motif. Moreover, the activation of autophagy during ER stress seems to be enhanced by calreticulin overexpression, and to alleviates ER stress. In addition, there is direct evidence that calreticulin, by translocating to the cell surface upon ER stress, can interact directly with the HLA-shared epitope and activate an innate immune response ^38,39^.

Finally, while our experimental set up did not address the citrullination process, we believe that native epitopes from described autoantigens could contribute to epitope spreading during the course of RA. Interestingly the seven autophagy related autoantigens that we have identified have been described to be targeted by ACPA. Since citrullination of these autoantigens have been precisely characterized, in a canonical study defining the citrullinome cellular atlas ^40^, we hypothesize that CD4 T cell response that sustain ACPA-producing B cell response is not necessarily directed against citrullinated epitopes. This hypothesis might be particularly interesting in the context of tertiary lymphoid structures. In this context, local stromal antigens that are expressed in synovial fibroblast can be presented through autophagy to CD4 T cells.

We propose that proteins from the ER stress response become targets of the immune response as autoantigens and contribute to epitope spreading during RA. In this context the interplay between ER stress and autophagy could contribute to the break of peripheral tolerance, with autophagosomes delivering stress proteins to the MHC class II compartment.

**In conclusion,** the study of a monoallelic DRB1*04:01 repertoire, strongly associated with the risk of developing RA, has unraveled autophagy as the pathway responsible for the processing of more than 10 autoantigens. Interestingly, in non-professional APCs, such as synovial fibroblasts that are key players in the pathogenesis of RA, autophagy is upregulated and contribute to the MHC II processing of 4 described RA autoantigens. Our findings describe a new mechanism of how autophagy can contribute to autoimmune disorders.

## Authors Contributions

N. Madelon Design Conduction and Analysis of most of the experiments, Writing -review & editing; M. Stumpe: Design, Conduction and Analysis of mass spectrometry experiments; A. Notto: Analysis of T cell activation experiments. J.Racle: Analysis of peptidomic data Writing -review & editing; E. Saiji: Histological analysis. N.Yeremenko: Provided with human RA samples. Conceptualization, and Discussion. D.Gfeller : Analysis of peptidomic data Writing -review & editing C. Ospelt: Conceptualization, Discussion and Writing -review & editing; J. Dengjel: Conceptualization, Methodology, Writing -review & editing; M. Gannagé: Conceptualization, Project administration, Supervision, Resources, Writing -original draft, Writing -review & editing.

## Acknowledgements

We would like to thank Pr Cem Gabay, Pr Didier Hannouche (University of Geneva), Pr Steffen Gay (University of Zurich), and Pr Dominique Baeten (University of Amsrerdam) for their help in providing access to RA and OA patient samples. We would like to thank Alexandre Ghounaris, for his contribution to the qPCR experimental set up of the synovial biopsies.

Monique Gannagé is funded by the Swiss national fund (310030_169938), and (310030_200677), the Institute of Rheumatology Research (Switzerland), the Novartis foundation (19C194) and the Fondation Pierre Mercier pour la Science

## Materiel and Methods

### Contact for reagent and resource sharing

Further information and requests for resources and reagents should be directed to and will be fulfilled by the lead contact, Monique Gannagé

### Cells

#### HLA-DRB1*04:01^+^ synovial fibroblasts and PBMCs of RA patients

HLA-DRB1*04:01^+^ synovial fibroblasts were obtained from synovial tissues of RA patients undergoing joint replacement surgery at the Schulthess Clinic, Zurich, Switzerland. Ethic permission was granted by the ethic commission of Zurich and informed consent was obtained. All patients fulfilled the revised criteria for RA ^41^. Synovial tissues were digested with dispase (37°C, 1h) and cultured by adherence at low passages in DMEM supplemented with 10% fetal calf serum, 1% penicillin-streptomycin; 2mM L-glutamine, 10mM HEPES and 0.2% amphotericin B (Gibco). EDTA blood was collected from the same patients undergoing joint replacement surgery and PBMCs were isolated by Ficoll-Paque Plus (VWR).

#### Synovial tissue specimens

Synovial tissue specimens were obtained from healthy donors (HD, from RA and OA patients either from ultrasound guided biopsies or during joint replacement surgery: (Department of Rheumatology and Department of Orthopedic Surgery, Zurich, Switzerland) (Department of Rheumatology and Department of Orthopedic Surgery, Geneva, Switzerland), and Department of Rheumatology, Amsterdam, Netherlands). Patients fulfilled the American College of Rheumatology criteria for classification of RA ^41^ or OA ^42^. All patients provided informed consent according to legal procedures from the local hospital ethical board.

#### 293T expressing HLA-DRB1*04:01 (HLA-DR4 cells)

293T cells stably transfected with cDNA encoding HLA-DRB1*04:01 (a gift from Pr Munz, Zurich) were cultured by adherence in DMEM supplemented with 10% fetal calf serum and 1% penicillin-streptomycin.

### Method details

#### CRISPR/Cas9 mediated gene disruption of Atg12 in 293T-DR4 cells

Specific short guide RNA (sgRNA) for Atg12 genes were designed using the genome engineering tool provided online by the Zhang laboratory (http://tools.genome-engineering.org). The following sgRNA sequences were used:

ATG12 (1) AAACACTTCAATTGCTGCTGGAGGC
ATG12 (2): AAACAGGCACTACTAAAAGGGGCAC

Oligonucleotide pairs were flanked with BsmBI restriction sites and cloned into the pSpCas9(BB)-2A-GFP(PX458) plasmid (gift from Pr Schmolke). The ligation step was performed using the rapid DNA ligation kit (ThermoFischer) according to the manufacturer’s instructions. The different CRISPR/Cas9 constructs obtained after ligation were analyzed by sequencing (Microsynth).

293T-DR4 cells were transfected with the CRISPR/Cas9 constructs by using lipofectamine 2000 (ThermoFischer). Briefly, 0.25×10^6^ 293T-DR4 cells were transfected with 4µg of plasmid in 6 well plates. After 48h, GFP^+^ cells were single-cell sorted by flow cytometry in 96 well plates, on a Beckman Coulter MoFlow Astrios. Individual clones were expanded and screened by western blot for LC3 and Atg5-12 expression. We used for all our experiments a selected clone of 293T-DR4-Atg12^−/-^ cell line.

#### Cocultures of HLA-DRB1*04:01^+^ synovial fibroblasts with matched autologous PBMC

HLA-DRB1*04:01^+^ synovial fibroblasts were cultured with 200U/ml of IFNg in normal complete medium or in starvation medium for 48h. Synovial fibroblasts were cocultured with matched autologous PBMC for 48h, at 3 different effectors to targets ratio: 10/1, 5/1 and 2/1. Of note all ratios could not be run for all samples due to the limited amount of available samples.

#### MHC class II immunoprecipitation

The first step was antibody coupling to CNBr sepharose beads. CNBr sepharose beads (GE healthcare) were pre-activated in a 1mM HCl buffer, and washed 5 times in a 100mM NaHCO_3_, 500mM NaCl coupling buffer (pH8.3). CNBr sepharose beads were then coupled to anti-human HLA-DR (L243) antibodies overnight at 4°C under rotation in coupling buffer. To remove uncoupled antibodies, beads were washed with coupling buffer and the reaction was quenched with 100mM Tris-HCl. Beads were finally washed alternatively in a 100mM Tris-HCl, 500mM NaCl buffer (pH8) and in a 100mM NaOAc, 500mM NaCl buffer (pH4).

Cells used for immunoprecipitation (293T-DR4-Atg12^+/+^ or 293T-DR4-Atg12^−/-^) or INFgamma treated synovial fibroblasts) were washed in PBS and lysed in a buffer of 1% CHAPS, 20mM TrisHCl (pH8), 5mM EDTA, 0.04% NaN3, 1mM PMSF, 100µM iodoacetamide buffer supplemented with proteases inhibitors (Roche) for 1h at 4°C under agitation. Lysates were obtained after centrifugation at 15000rpm for 20 minutes and clarified twice at 3000rpm for 20 minutes. Equal protein amounts from lysates were used after quantification by BCA protein assay kit (Pierce). Lysates were further immunoprecipitated overnight at 4°C under rotation using CNBr sepharose beads coupled to either anti-mouse IA-IE or anti-human HLA-DR antibodies. The following day, beads were washed 3 times with lysis buffer, 6 times in a 50mM TrisHCl 250mM NaCl buffer (pH8), and 6 times in a 50mM NaCl buffer (pH8). MHC-II-peptide complexes were eluted from beads by incubating them with 10% acetic acid for 10 minutes at room temperature and were than stored at −80°C before mass spectrometry analysis. A small fraction of the samples was washed in PBS, ultra-centrifugated and loaded on SDS-polyacrylamide gel to check for MHC-II immunoprecipitation.

### LC-MS/MS analyses

LC-MS/MS measurements were performed on a QExactive (QE) HF-X mass spectrometer coupled to an EasyLC 1200 nanoflow-HPLC (all Thermo Scientific). MHC peptides were fractionated on a fused silica HPLC-column tip (I.D. 75 μm, New Objective, self-packed with ReproSil-Pur 120 C18-AQ, 1.9 μm (Dr. Maisch) to a length of 20 cm) using a gradient of A (0.1% formic acid in water) and B (0.1% formic acid in 80% acetonitrile in water): samples were loaded with 0% B with a flow rate of 600 nL/min; peptides were separated by 5%–30% B within 85 min with a flow rate of 250 nL/min. Spray voltage was set to 2.3 kV and the ion-transfer tube temperature to 250°C; no sheath and auxiliary gas were used. The mass spectrometer was operated in the data-dependent mode; after each MS scan (mass range m/z = 370 – 1750; resolution 120’000) a maximum of twelve MS/MS scans were performed using a normalized collision energy of 25%, a target value of 5′000 and a resolution of 30’000. MS raw files were analyzed using MaxQuant (version 1.6.2.10) ^43^ using Uniprot full-length human (March, 2016) and mouse (April 2016) databases and common contaminants such as keratins and enzymes used for in-gel digestion as reference. Protein amino-terminal acetylation and oxidation of methionine were set as variable modifications. The MS/MS tolerance was set to 20 ppm using unspecific as enzyme specificity. Peptide and protein FDR based on a forward-reverse database were set to 0.01, minimum peptide length was set to 9, the maximum to 25, the minimum score for modified peptides was 40, and minimum number of peptides for identification of proteins was set to one, which must be unique. The “match-between-run” option was used with a time window of 1 min. MaxQuant results were analyzed using Perseus ^44^. Data are available via ProteomeXchange with identifier PXD020624; username, password: 0fk87Vsc ^45^.

### Methods for peptidomics analysis

#### HLA-DRB1*04:01 ligands analysis

To determine peptides that were presented by HLA-DRB1*04:01 in the MHC-II peptidomics data, we used MixMHC2pred-2.0 ^12^ the 293T cells and fibroblasts samples. Citrullinated arginine was replaced by the letter “X” when running MixMHC2pred. The peptides’ N-/C-terminal contexts used by MixMHC2pred were added by mapping the peptides to the human proteome and keeping the 3 residues upstream of the peptide + 3 N-terminal residues of the peptide (N-terminal context) and the 3 C-terminal residues of the peptide + 3 residues downstream of the peptide (C-terminal context).

Peptides were retained as HLA-DRB1*04:01 ligands when their %Rank returned by MixMHC2pred was better than 20, corresponding to a threshold of predicted weak binding. Peptides of length shorter than 12 amino acids or longer than 21 amino acids were not considered as these are not predicted by MixMHC2pred and are mostly contaminants. For the subsequent analyses, only these predicted HLA-DRB1*04:01 ligands were kept, the other peptides being either contaminants peptides or presented by other HLA-DR alleles in the heterozygote fibroblasts samples.

#### Analysis of proteins differentially presented

In the 293T cells samples, the MS peptide intensities of each HLA-DRB1*04:01 ligand was normalized by the sum of all HLA-DRB1*04:01 ligands from the given sample. The peptides observed in one condition (Wild-type (ATG12^+/+^)or ATG12^−/-^) but not the other from the same batch (i.e., batch 0407, 1007, …) were set to a value of 0. Peptides observed in less than 3 batches were removed. **Supplementary table 2** lists these values, multiplied by 100. We then performed a two-sided paired Wilcoxon signed-rank test of these peptides grouped by their source gene to determine proteins that were differentially presented between the WT and ATG12^−/-^ conditions. P-values were adjusted for multiple testing by Holm correction.

### Quantitative RT-PCR (qPCR)

Total RNA was extracted using the RNeasy mini kit according to the manufacturer’s instructions (QIAGEN). Reverse transcription was performed using superscript II. The RT product was diluted and used as a template for quantitative PCR on a CFX Connect real-time detection system (Bio-Rad) using Sybr Green (Kapa Biosystems). The following primers (Microsynth) were used:

LC3 forward 5’-AGACCTTCAAGCAGCGCCG-3’
LC3 reverse 5’-ACACTGACAATTTCATCCCG-3’
Atg5 forward 5’-CAACTTGTTTCACGCTATATCAGG-3’
Atg5 reverse 5’-CACTTTGTCAGTTACCAACGTCA-3’
Atg12 forward 5’-TCTTCCGCTGCAGTTTCC-3’
Atg12 reverse 5’-GGAGCAAAGGACTGATTCACA-3’

### Histology

Paraffin-embedded synovial biopsies of osteoarthritis (OA) and rheumatoid arthritis (RA) patients were stained for LC3, p62, MHC-II and vimentin proteins. Sections were permeabilized with 0.1% triton and blocked in a 1%BSA 2% FCS buffer for 30 minutes. Slides were incubated with the following primary antibodies: mouse anti-LC3 (Nanotools), rabbit anti-p62 (MBL international), mouse anti-HLA-DR-DP-DQ (Abcam) and rabbit anti-vimentin (Cell Signaling Technology) overnight at 4°C. Slides were further incubated with the following secondary antibodies: Alexa fluor488-conjugated anti-mouse and Alexa fluor555-conjugated anti rabbit IgG (ThermoFischer) for 1h at room temperature. Nuclei were visualized using prolong gold antifade mounting medium with DAPI (ThermoFischer). Z-stacks were performed on a LSM800 confocal microscope (Zeiss). The number of autophagosomes per cell and the mean fluorescent intensity of cytoplasmic p62 were evaluated using the Imaris software (version 9.3.1).

### Western blot

Cells were lysed in a 1% NP-40 buffer supplemented with protease inhibitors (Roche). Equal protein amounts from lysates were used after quantification by BCA protein assay kit (Pierce). Protein samples were denatured by Laemmli buffer, separated by SDS-polyacrylamide gel electrophoresis and transferred to PVDF membranes. Immunoblots were blocked and incubated with primary antibodies in PBS 0.05% Tween-20 5% non-fat dry milk overnight at 4°C. The following primary antibodies were used: mouse anti-Atg5-12 (Nanotools), and mouse anti-vinculin (Sigma); and the following secondary antibodies: HRP-conjugated anti-mouse or anti-rabbit IgG (Bio-Rad).

## QUANTIFICATION AND STATISTICAL ANALYSIS

Statistical parameters including the exact value of n, the dispersion, the precision of measures (mean +/- SEM) and the statistical significance are reported in the figures and figure legends. Statistical analysis was performed in GraphPad Prism software (version 8.0.2). Student’s t test or Mann Whitney unpaired non-parametric t test was used (unless otherwise stated in the figure legend). The statistical significance level was set as p values and depicted in figures as following: *p < 0.05, **p < 0.01, ***p < 0.001.

