## supplemental Materials for "Autophagy shapes the peptide repertoire of rheumatoid arthritis-associated HLA class II alleles"

**Figure S1 (Related to Figure 1) (A)** Western blot showing Atg5-12 expression in Atg12$^{+/+}$ 293T-DR4 and Atg12$^{-/-}$-293T-DR4 cell lines. **(B)** Histogram showing flow cytometry analysis of HLA-DR expression in Atg12$^{-/-}$ and Atg12$^{+/+}$293T-DR4 cells. Bar graphs indicate the relative fold change of the MFI of HLA-DR expression in Atg12$^{-/-}$ cells, compared to Atg12$^{+/+}$ cells. Data are from 3 independent experiments. **(C)** Motifs describing binding specificities were obtained by running MoDec on the MHC class II peptidomics data from each sample separately. Peptides of length 9 and longer were considered. In each sample, a single motif corresponded to non-contaminants peptides. The corresponding frequency of each amino acid at the 9 binding core positions of these motifs were used to perform a hierarchical clustering based on Ward.D2 method, showing that samples did not cluster based on their ATG12 status. **(D)** Pie charts showing the number and relative frequency (in %) of peptides that are enriched (pink) in the HLA-DR peptidome of autophagy deficient 293-T DR4 cell line (Atg12$^{-/-}$) as compared to (Atg12$^{+/+}$) paired sample. The gray repertoire represents the pool of peptides that are stable between both cell lines. One representative paired experiment is shown out of five **(E)** Graphs showing the normalized intensity (in % of the repertoire) of every peptide from the related HLA class 1 molecules that are enriched in (Atg12$^{-/-}$) cells as compared to wt. **(F)** Heat Map representing the relative intensity of all proteins that are significantly enriched in MHC Class II peptides eluates from Atg12$^{-/-}$ vs Atg12$^{+/+}$ -DRB1*04:01 cells. A Z normalization of the MS intensity was done for every antigen, using R software.

**Figure S2 (related to Figure 2 and 3) (A)** Immunofluorescence analysis of p62 expression in synovial biopsies of OA and RA patients. Confocal images of representative sections from OA and RA biopsies: p62(red), LC3(green), Dapi(blue). Scale bar: 10um. Bar graphs represent the mean MFI of cytoplasmic p62 in OA and RA biopsies. Nuclear P62 staining was excluded from the analysis. **(B)** Representative histogram showing HLA-DR expression in synovial fibroblast, treated with INF gamma for 48h, in normal medium or under starvation conditioned medium.

(C) Western blot showing LC3 expression in synovial fibroblast under different conditions: starvation-medium (48h), INFgamma or CQ (50uM) treatment. Vinculin expression is used as a loading control. (D) Bar graph represent the relative expression of LC3-II in synovial fibroblasts under the depicted conditions.

**Suppl. Table 1**

**List of peptides that were eluted from HLA-DR4 Atg12$^{-/-}$ and control wild-type cells from 5 independent experiments.** Experiments are numbered as 0407, 293T, 1004, 2004, and 2703. The N- and C-terminal context of each peptide is also indicated, as well as the predicted %Rank obtained from MixMHC2pred-2.0 (values are left empty for peptides shorter than 12 amino acids or longer than 21 amino acids; see *Methods*). Peptides with a %Rank < 20 were annotated as HLA-DR4 ligands.

**Suppl. Table 2**

**Relative frequencies of HLA-DRB1*04:01 ligands from 5 independent experiments of peptides eluates from DR4 ATG12$^{-/-}$ and wild-type control paired cell lines.** Only peptides from Supplementary Table 1 predicted to be HLA-DR4 ligands were considered. The MS peptide intensity from each ligand was normalized by the sum of all HLA-DRB1*04:01 ligands from the given sample and multiplied by 100 in this table (columns *ATG12* and *WT*). Ligands observed in an experiment in only one condition (ATG12$^{-/-}$ or wild-type) were set to a value of 0 in the other condition.

**Suppl. Table 3**

**List of peptides that were eluted from 3 synovial fibroblast lines SKH100, SKH108, and SKH124.** The predicted %Rank score from MixMHC2pred-2.0 towards HLA-DRB1*04:01 is indicated (see *Methods*); peptides indicated as "FAUX/FALSE" in the *DR4_ligand* column could be either contaminant peptides or could be presented by other HLA-DR alleles in the heterozygote fibroblasts samples.

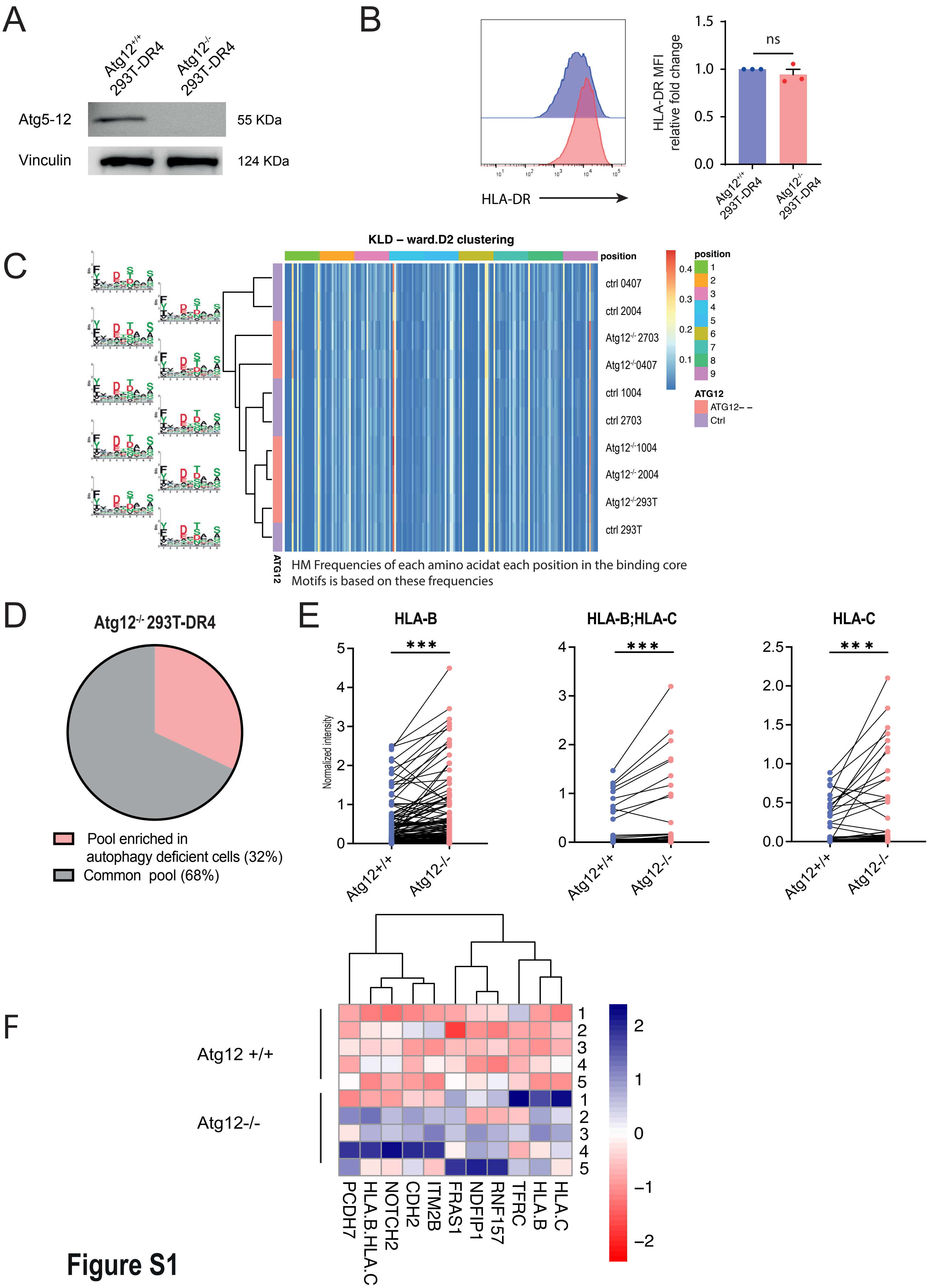

A
Atg12+/+ 293T-DR4
Atg12-/- 293T-DR4
Atg5-12
Vinculin
55 KDa
124 KDa
B
HLA-DR
HLA-DR MFI relative fold change
ns
Atg12+/+ 293T-DR4
Atg12-/- 293T-DR4
C
KLD – ward.D2 clustering
position
ctrl 0407
ctrl 2004
Atg12-/- 2703
Atg12-/- 0407
ctrl 1004
ctrl 2703
Atg12-/- 1004
Atg12-/- 2004
Atg12-/- 293T
ctrl 293T
ATG12
ATG12– –
Ctrl
HM Frequencies of each amino acidat each position in the binding core
Motifs is based on these frequencies
D
Atg12-/- 293T-DR4
Pool enriched in autophagy deficient cells (32%)
Common pool (68%)
E
HLA-B
HLA-B;HLA-C
HLA-C
Normalized intensity
Atg12+/+
Atg12-/-
F
Atg12 +/+
Atg12-/-
PCDH7
HLA.B.HLA.C
NOTCH2
CDH2
ITM2B
FRAS1
NDFIP1
RNF157
TFRC
HLA.B
HLA.C


**Figure S1**

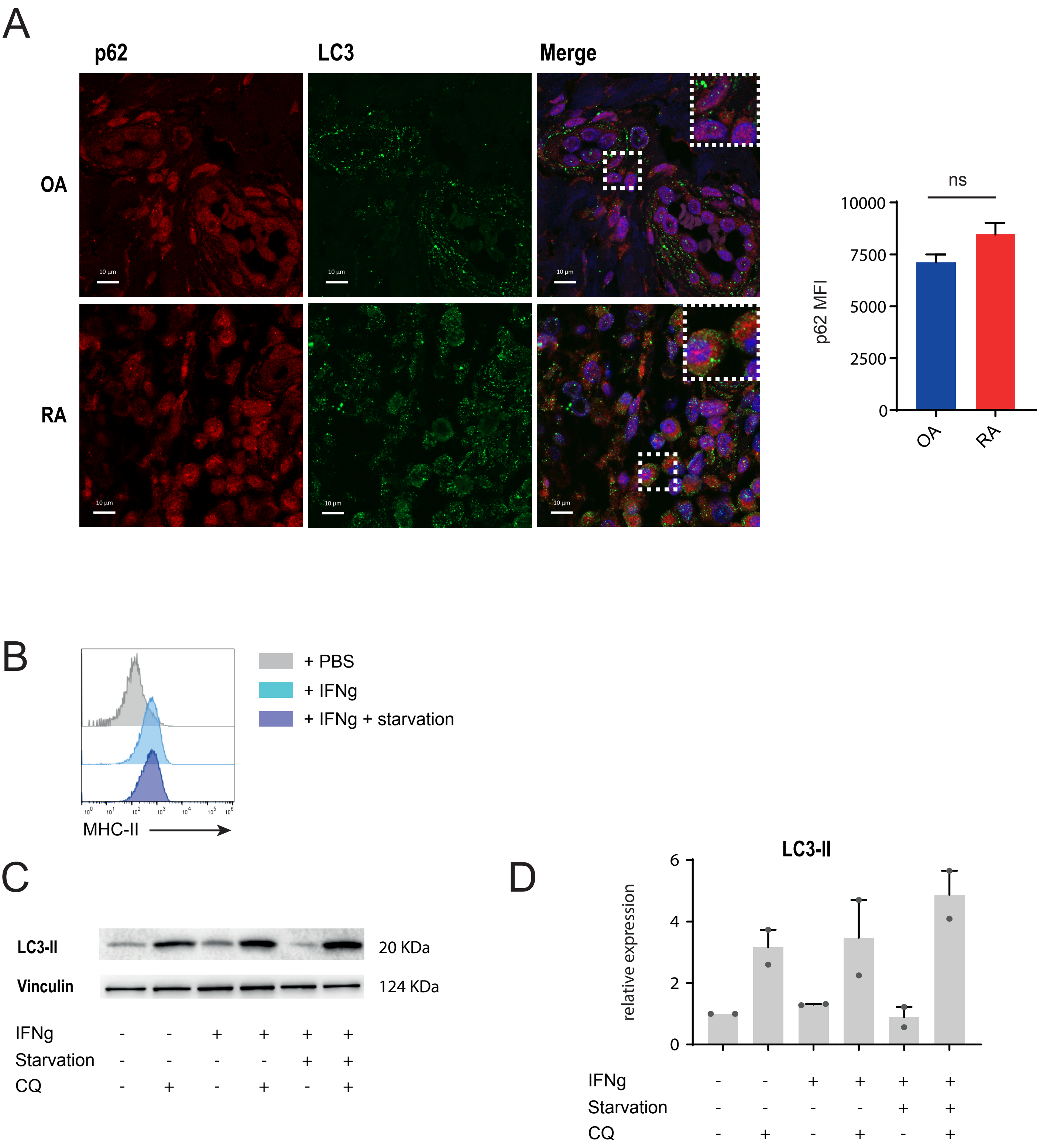
A
p62
LC3
Merge
OA
RA
10 µm
ns
10000
7500
5000
2500
0
p62 MFI
OA
RA
B
+ PBS
+ IFNg
+ IFNg + starvation
MHC-II
C
LC3-II
Vinculin
20 KDa
124 KDa
IFNg - - + + + +
Starvation - - - - + +
CQ - + - + - +
D
LC3-II
relative expression
6
4
2
0
IFNg - - + + + +
Starvation - - - - + +
CQ - + - + - +

**Figure S2**
